# ALPINE: A Scalable Pipeline for Comprehensive Classification of Gene-Editing Outcomes from Long-Read Amplicon Sequencing

**DOI:** 10.64898/2026.03.27.714831

**Authors:** Yu Chen, Xing-Huang Gao, Athea Vichas, Jianbin Wang, Ryan Golhar, Isaac Neuhaus

**Author notes:** Co-corresponding authors: Yu Chen; Xing-Huang Gao. Co-first authors; these authors contributed equally to this work (Yu Chen, Xing-Huang Gao).

## Abstract

CRISPR genome editing has enabled precise genetic modification for gene and cell therapies, but edits often produce heterogeneous on-target outcomes, including homology-directed repair (HDR) knock-ins, DNA repair template integrations, and structural variants. Existing tools are frequently limited to short reads or lack viral vector-specific integration categories needed for therapeutic development. Here, we present ALPINE (Amplicon Long-read Pipeline for INtegration Evaluation), a scalable and reproducible pipeline for classifying and quantifying gene-editing outcomes from long-read amplicon sequencing. ALPINE classifies reads into 10+ categories, including DNA repair vector integration subtypes, and performs variant calling near the gene-edited site with batch, multi-sample reporting. Uniquely, ALPINE can distinguish between cells treated with multiple DNA repair vectors and identify distinct molecular features, such as inverted terminal repeats (ITRs), enabling comprehensive characterization of complex gene editing outcomes. Benchmarking on simulated datasets showed high accuracy, and application to edited T cell samples demonstrated comprehensive gene-editing outcome profiling. Supplementary data are available online.

**Availability:** ALPINE is implemented in Python and distributed as Docker containers with Common Workflow Language (CWL) support for cloud deployment. The pipeline is available under MIT license at https://github.com/Maggi-Chen/ALPINE.

## 1. Introduction

CRISPR-Cas genome editing has revolutionized cell-based therapies by enabling precise genetic modifications, such as gene knockouts and transgene knock-ins through homology-directed repair (HDR) with viral vector donors (Doudna and Charpentier, 2014). However, repair of CRISPR-induced double-strand breaks can yield heterogeneous outcomes, including structural variants and viral vector integrations. While recent studies have illustrated that in addition to intended perfect HDR, gene editing can result in additional repair outcomes, including large deletions and DNA repair template integration (Hunt et al., 2023). In adeno-associated virus (AAV)-mediated editing, approximately 1–2% of reads contain AAV integrations, including full-length AAV genomes and fragments with inverted terminal repeat (ITR) sequences (Simpson et al., 2023). Such unintended modifications at gene-edited loci (Figure S1) can disrupt transgene expression, compromise efficacy, or introduce safety risks, making accurate characterization critical for regulatory evaluation of gene-edited therapies. Despite the importance of capturing gene-editing outcomes, existing tools have major limitations for long-read amplicon data from Cas-mediated editing. CRISPResso2, one of the most widely used genome editing analysis tools, is designed for short-read Illumina data with maximum practical read length at ∼600 bp, limiting detection of large structural variants, full-length HDR knock-ins, and multi-kilobase AAV integrations (Clement et al., 2019). The knock-knock pipeline supports long reads and profiles HDR outcomes (Canaj et al., 2019), but lacks AAV-specific categories (e.g., integration with/without ITR) and supports only a single homologous donor template per target site, preventing attribution when multiple AAV vectors may integrate at the same locus. Its multi-aligner workflow (BLASTn, STAR, minimap2) also increases computational overhead (Canaj et al., 2019). As a result, researchers often rely on manual counting of AAV integrations from long-read alignments (Simpson et al., 2023).

To address these limitations, we developed ALPINE (Amplicon Long-read Pipeline for INtegration Evaluation), a scalable bioinformatics pipeline for automated classification and quantification of gene-editing outcomes from long-read amplicon sequencing data (PacBio HiFi). ALPINE classifies reads into 10+ variant categories, including HDR knock-ins and AAV vector insertions with or without ITR sequences, and supports multiple AAV vectors to identify which vector contributed each integration event. ALPINE also performs variant calling within a defined window around the cleavage site. Built with Docker containers and CWL workflows, ALPINE enables reproducible, cloud-deployed analysis suited to high-throughput studies and regulatory environments.

## 2. Methods

### 2.1 Workflow overview

The ALPINE pipeline uses a multi-cloud launcher framework to execute a per-sample analysis workflow and merge per-sample outputs into a final summary table. Each workflow consists of five sequential steps: read filtering, alignment, classification, counting, and merging (Figure 1A). The inputs include FASTQ files, reference sequences (wild-type, HDR, and optionally AAV integrations), a configuration file specifying cleavage site coordinates and vector element boundaries (ITRs and transgene), primer and homology arm sequences, and optional quality thresholds. Outputs include per-sample classification tables, quality control reports, and a merged summary table.

**Figure 1.**
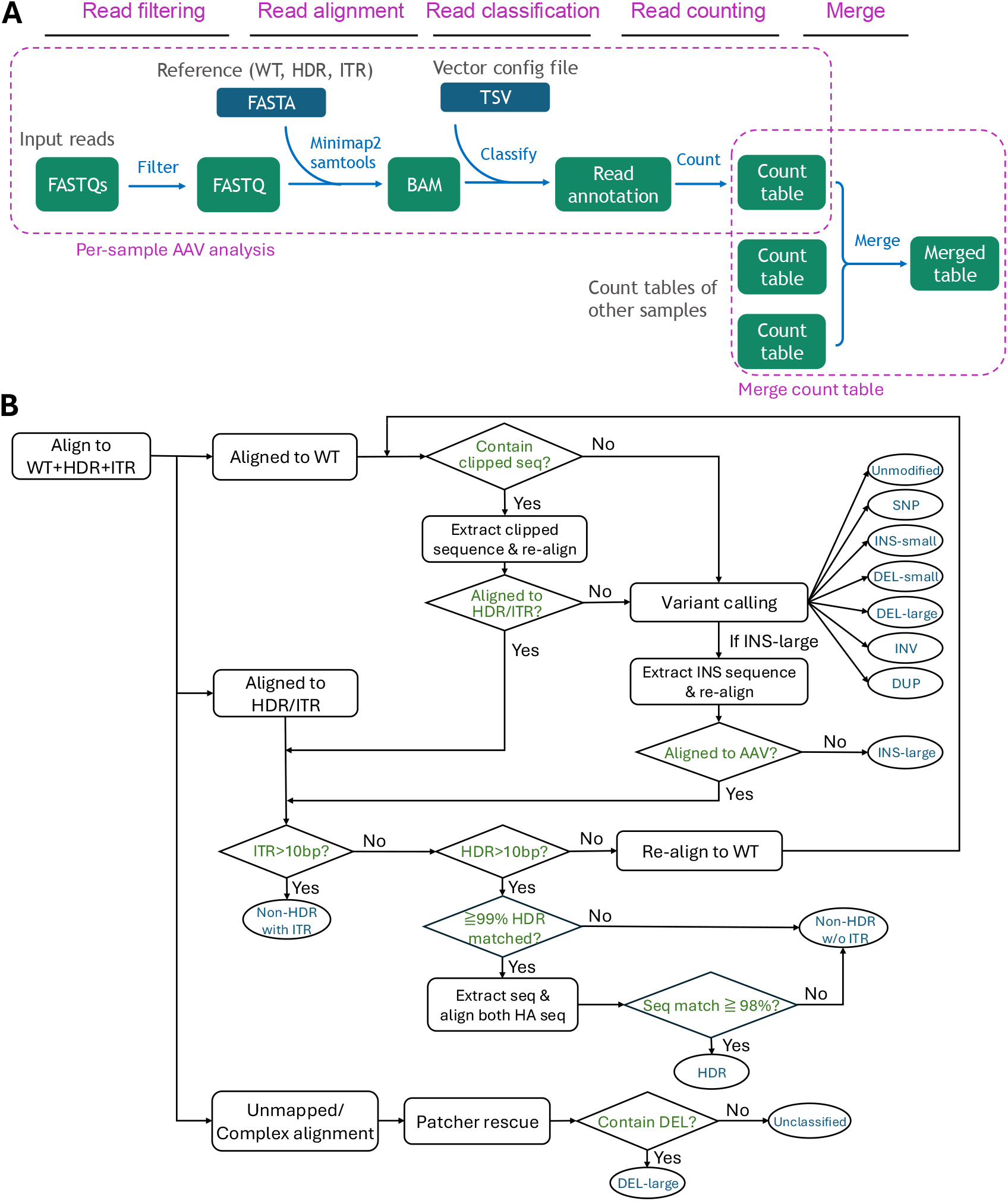
Overview of the ALPINE pipeline for long-read amplicon sequencing analysis. **(A)** ALPINE workflow includes five major steps: read filtering, alignment with minimap2, read classification, read counting, and table merging. **(B)** Read classification algorithm flowchart showing the decision logic used to assign reads to variant categories based on alignment to wild-type, HDR, and AAV integration reference sequences, including three re-alignment modules and the patcher rescue step.

### 2.2 Read filtering and alignment

Reads are first filtered to ensure completeness and sequencing quality. Primer sequences are used to search for forward and reverse primers in the first and last 100 bp of each read. Reads containing both primers are retained. Read average sequencing quality is calculated and reads below the threshold (default Q30) are removed. A quality control report with histogram plots of read length and quality distributions before and after filtering is generated (Figure S2).

Filtered reads are aligned to reference sequences using minimap2 with the map-hifi preset optimized for PacBio HiFi data (Li, 2018). The reference FASTA includes WT sequence, HDR knock-in sequence, and AAV vector integration sequences. Additional references can be included to capture other expected editing outcomes (e.g., antisense transgene knock-in). The config file provides 1-based coordinates for WT cleavage site, HDR transgene boundaries, and ITR boundaries in the AAV integration reference. Aligned reads are sorted and indexed using SAMtools (Li et al., 2009).

### 2.3 Read classification

Reads are classified based on alignment results (Figure 1B). Classification involves a series of alignment and re-alignment steps:

- **Reads aligned to WT reference**: Variant calling is performed within ±20 bp of the cleavage site, assigning each read to one of the variant types: unmodified, DEL-large (≥50bp), DEL-small (<50bp), INS-large (≥50bp), INS-small (<50bp), unmodified-with-SNP, INV (inversion), DUP (duplication). Large insertions are extracted and re-aligned to all references to determine whether inserted sequences originate from transgene or DNA repair template (e.g. AAV vector).
- **Reads aligned to HDR/AAV integration references**: Reads aligned to HDR or AAV references are evaluated for transgene and AAV content to distinguish between perfect HDR knock-in and AAV integration (referred as NonHDR). AAV insertions are further classified into NonHDR-with-ITR and NonHDR-without-ITR categories based on ITR sequence presence.
- **Re-alignment modules**: Three re-alignment modules address imperfect initial alignments. (1) Inserted-sequence re-alignment for WT-aligned reads with large insertions (≥50bp), extracting inserted sequences and re-aligning to HDR/AAV references. (2) Clipped-sequence re-alignment for WT-aligned reads with unmapped clipped segments (≥100bp), extracting clipped sequences and re-aligning to check for transgene content and call variants. (3) WT re-alignment for reads aligned to HDR/AAV references that lack transgene sequence, re-aligning full read to WT reference for variant calling.
- **False-negative rescue**: A “patcher” module re-analyzes unclassified reads by realigning them to the WT reference using minimap2 v2.16 with map-pb preset, allowing detection of large deletions missed in initial classification. Reads matching large deletion pattern are rescued and reassigned to DEL-large.

Additional details on the variant classification methods are provided in Supplementary Note 1.

### 2.4 Read counting and merging

Following read classification, ALPINE quantifies the number of reads within each variant category for each sample, and outputs the results as a count table. A pie chart is generated to visualize category proportions (Figure S3). When multiple samples are processed, count tables are merged into a combined summary table (Figure S4). For experiments involving different target sites or transgenes, the merged table includes all vector-specific categories with zero filling for categories not applicable to a sample.

### 2.5 Cloud integration and scalability

ALPINE is implemented using CWL and Docker containers for platform agnostic deployment. Each step runs in an isolated Docker container. A multi-platform launcher manages platform-specific configuration and resource allocation, supporting SevenBridges Genomics, Amazon HealthOmics and Arvados. This architecture enables scalable analysis of large datasets while maintaining reproducibility required for regulatory submissions.

## 3. Results

### 3.1 Benchmark on simulated datasets

We evaluated the accuracy of ALPINE’s read classification using simulated datasets representing a diverse range of gene editing outcomes. Fifteen groups of simulated reads were generated using PBSIM3 (Ono et al., 2022), with each group containing 20,000 reads corresponding to a specific editing outcome (Supplementary Note 2). These outcomes included HDR knock-in, non-HDR vector integration with full ITR sequences (forward and reverse complement), unmodified, one-sided ITR retention (left and right, forward and reverse), two-sided ITR with 50% deletion (forward and reverse), one-sided ITR with 50% deletion (left and right), and vector integration events containing SNPs within ITR regions, encompassing both-sided, left-sided, and right-sided ITR configurations.

Using default parameters, ALPINE achieved 100.00% accuracy in 14 of the 15 simulated groups (Table S1). For the WT unmodified group, ALPINE correctly classified 97.60% (19,519/20,000) of reads as “Unmodified”, with the remaining reads reported as small indels (472) or SNPs (9), consistent with simulated sequencing errors near the cut site. Overall, these results demonstrate ALPINE’s ability to accurately classify gene-editing outcomes across a wide range of variant categories, while highlighting the value of stringent quality filtering for robust quantification.

### 3.2 Application to gene-edited T cell samples

We applied ALPINE to PacBio HiFi long-read sequencing data from five edited human T cell samples, each edited at two distinct genomic loci using corresponding CRISPR RNPs and two AAV-delivered transgene constructs, generating 10 datasets in total. Quality control assessment indicated high sequencing quality, with high average base quality scores and expected read-length distributions across all datasets. For each dataset, ALPINE produced per-sample outcome tables summarizing the distribution of reads across HDR, WT-derived outcomes (unmodified and indels), structural variants, and non-HDR integration categories (with/without ITR), along with standardized merged summaries across samples. These tabular outputs provide a compact view of editing efficiency and byproduct formation (unintended editing events), while the accompanying read-length and quality summaries support interpretation of dominant editing modes (e.g., knock-in versus knockout) and facilitate rapid cross-sample comparisons.

ALPINE successfully classified reads for all samples at both target sites (Tables S2–S3). Consistent patterns were observed across datasets: HDR knock-in events were predominant, while structural variants and unintended integration events (NonHDR with/without ITR) were detected at lower frequencies. Non-HDR AAV integrations and other unintended modifications represented a minor fraction of reads at both target sites (Table S4). Read-length distribution patterns for all five samples correlated strongly with ALPINE’s quantitative classification results, with samples showing higher knock-in frequencies displaying higher peaks in the knock-in–associated read-length range, and those with higher knockout frequencies showing correspondingly enhanced knockout peaks (Figure S5). This concordance between read-length distributions and ALPINE’s algorithmic classifications supports the accuracy of the pipeline’s gene-editing outcome quantification.

## 4. Discussion and Conclusions

Compared with existing tools, ALPINE provides automated, end-to-end quantification of on-target outcomes from long-read amplicon sequencing and is configurable for both viral and non-viral donor methods. ALPINE classifies reads into “donor-aware” categories (including AAV integration subtypes when ITR/transgene boundaries are provided) and supports multi-vector experiments by attributing each integration event to the corresponding vector. It combines fast minimap2 alignment with targeted re-alignment for ambiguous subsets, enabling efficient batch processing with standardized outputs suitable for high-throughput and regulatory settings.

In addition to regular read alignment, ALPINE includes several re-alignment modules to mitigate misclassification caused by imperfect alignments. For WT-aligned reads with large insertions, ALPINE extracts the inserted sequence and re-aligns it to HDR/AAV references; for reads with large soft-clipped segments, ALPINE re-aligns clipped sequences to recover transgene/vector content. In addition, a “patcher” module re-analyzes initially unclassified reads using a secondary alignment pass with minimap2 to recover large deletions missed in the initial classification.

One limitation of ALPINE is its current focus on PacBio HiFi data. Nanopore long read support is not yet implemented but could be added by adjusting minimap2 presets and downstream filtering thresholds. Additionally, ALPINE is designed for on-target amplicon sequencing and does not assess off-target editing or genome-wide AAV integration. Future work will explore integration with off-target detection methods and expansion of classification categories based on emerging gene editing modalities.

In conclusion, ALPINE provides an automated, cloud-deployable pipeline for classification and quantification of gene-editing outcomes from long-read amplicon sequencing data, including detailed characterization of AAV integration events. ALPINE addresses a key gap in automated analysis of long-read AAV integration outcomes and enables vector-specific and regulatory relevant assessment of complex editing outcomes for gene and cell therapy research.

## References

Doudna JA, Charpentier E. Genome editing: the new frontier of genome engineering with CRISPR-Cas9. Science. 2014.

Hunt, J.M.T., Samson, C.A., Rand, A.d. et al. Unintended CRISPR-Cas9 editing outcomes: a review of the detection and prevalence of structural variants generated by gene-editing in human cells. Hum Genet. 2023.

Simpson BP, Yrigollen CM, Izda A, Davidson BL. Targeted long-read sequencing captures CRISPR editing and AAV integration outcomes in brain. Molecular Therapy. 2023.

Clement K, Rees H, Canver MC, et al. CRISPResso2 provides accurate and rapid genome editing sequence analysis. Nature Biotechnology. 2019.

Canaj H, Hussmann JA, Li H, et al. Deep profiling reveals substantial heterogeneity of integration outcomes in CRISPR knock-in experiments. bioRxiv. 2019.

Li H. Minimap2: pairwise alignment for nucleotide sequences. Bioinformatics. 2018.

Li H, Handsaker B, Wysoker A, Fennell T, Ruan J, Homer N, Marth G, Abecasis G, Durbin R; 1000 Genome Project Data Processing Subgroup. The Sequence Alignment/Map format and SAMtools. Bioinformatics. 2009.

Ono Y, Hamada M, Asai K. PBSIM3: a simulator for all types of PacBio and ONT long reads. NAR Genom Bioinform. 2022.

