## Supplemental notes, tables, and figures for "ALPINE: A Scalable Pipeline for Comprehensive Classification of Gene-Editing Outcomes from Long-Read Amplicon Sequencing"

### Supplementary Information

#### Supplementary Note 1: ALPINE Read Classification Methods

ALPINE employs a multi-step classification algorithm that assigns each sequencing read to a variant category based on its alignment to reference sequences. The algorithm includes three re-alignment modules to handle imperfect initial read alignments, ensuring comprehensive classification of diverse editing outcomes.

**Module 1: Insertion re-alignment.** For reads initially aligned to the wild-type (WT) reference that contain insertions ≥50 bp near the cleavage site (±20 bp), ALPINE extracts the inserted sequence and re-aligns it against all reference sequences (WT, HDR, and AAV integrant). If the inserted segment aligns to an HDR or AAV integrant reference, the read is re-evaluated for transgene content and reclassified accordingly. Conversely, if the inserted sequence aligns only to the WT reference or fails to align to any alternative reference, the read is assigned to the INS-large category.

**Module 2: Unmapped/clipped sequence re-alignment.** For reads aligned to the WT reference that contain unmapped or soft-clipped sequences ≥100 bp, ALPINE extracts these sequences and re-aligns them to the HDR and AAV integrant reference sequences. This module identifies cases where part of a read correctly maps to the WT genome while another portion contains transgene or vector-derived sequence that was not captured by the initial whole-read alignment. Such mixed-origin reads indicate potential integration events that would otherwise be missed, enabling more comprehensive detection and classification of complex repair outcomes.

**Module 3: HDR/AAV integrant aligned read re-evaluation.** For reads initially aligned to HDR or AAV integrant reference sequences that lack detectable transgene sequence—based on the expected transgene boundaries defined in the configuration file—ALPINE re-aligns the entire read to the WT reference for variant calling. This module addresses cases in which a read is preferentially aligned to the HDR or ITR reference due to sequence homology within the arms but does not actually contain transgene sequence. By re-evaluating such reads against the WT reference, ALPINE prevents misclassification and ensures accurate assignment of true editing outcomes.

**Patcher module (Large-Del false-negative rescue).** After the primary classification step is complete, ALPINE applies a post-processing patcher module to recover potential false-negative large-deletion events. Reads initially categorized as Unclassified are re-examined by extracting their sequences directly from the BAM file and re-aligning them to the WT-only reference using minimap2 v2.16 with the *map-pb* preset. This specific minimap2 version is used because newer releases (≥2.28) tend to assign those reads as soft-clip instead of representing them in the CIGAR string, which can obscure true deletion events. The patcher detects large deletions (≥50 bp) using both CIGAR-based analysis within a ±20 bp window around the target site and segment-based analysis within a ±10 bp window. Only DEL-large reads are rescued to maintain conservative reclassification, and the read name classification file is updated in place with the corrected assignments.

**Multi-vector classification.** When multiple AAV vectors are used within the same experiment—such as distinct AAV vectors delivering different transgenes to separate genomic loci—ALPINE assigns each integration event to the correct vector based on the specific reference sequence to which the read aligns. This enables vector-resolved quantification of editing outcomes even in complex multiplexed designs. The resulting count table contains per-vector columns for HDR, Non-HDR-with-ITR, and Non-HDR-without-ITR categories (e.g., HDR-VectorA, Non-HDR-with-ITR-VectorA, Non-HDR-without-ITR-VectorB). These vector-specific measurements allow researchers to independently track editing efficiency, integration profiles, and ITR retention for each vector at its corresponding target site.

#### Supplementary Note 2: Simulated Dataset Generation

Simulated datasets were generated to benchmark ALPINE's classification accuracy across all major editing outcome categories. For each simulation group, a template sequence was constructed by computationally assembling the expected read structure from the reference sequences. PacBio HiFi-like sequencing errors were introduced at a rate consistent with the expected HiFi error profile (approximately 0.1–0.5% per-base error rate).

The 15 simulation groups were designed as follows:

• Group 1 (HDR): Full-length HDR knock-in sequence flanked by homology arms and genomic sequence.

• Groups 2–3 (Full AAV integrant): Complete AAV integration sequence in forward and reverse complement orientations.

• Group 4 (WT/Unmodified): Wild-type reference sequence without modifications.

• Groups 5–8 (One-sided AAV ITR region): AAV integration with ITR sequence on one side only (left/right, forward/reverse).

• Groups 9–10 (Two-sided ITR, 50% deletion): AAV integration with 50% of the ITR sequence deleted, in forward and reverse orientations.

• Groups 11–12 (One-sided ITR, 50% deletion): AAV integrant with partial ITR integration on one side, with 50% deletion of the ITR sequence.

• Groups 13–15 (ITR with SNPs): AAV integration with introduced SNPs at various positions in ITR regions (two-sided, one-sided right, one-sided left).

Each group contained 20,000 simulated reads. Reads were processed through the ALPINE pipeline with default parameters. Classification accuracy was calculated as the percentage of reads assigned to the expected category.

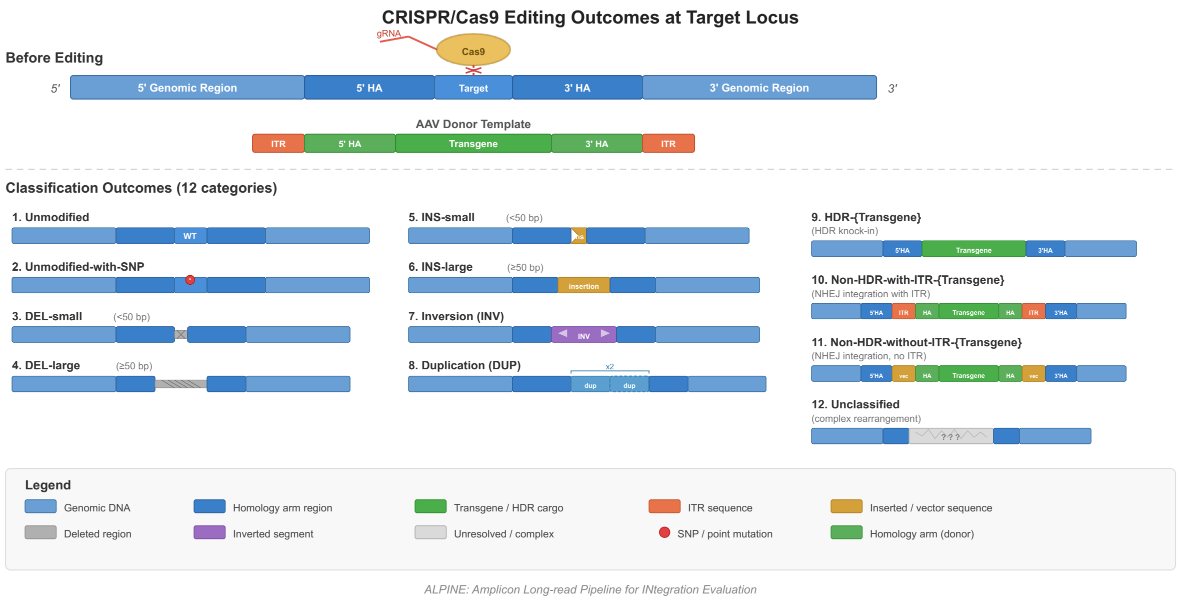


**Figure S1. Cartoon schematic of CRISPR/Cas9 editing.** Various genomic outcomes at the target site, including unmodified, unmodified-with-SNP, small and large deletions, small and large insertions, inversions, duplications, HDR knock-in, non-HDR vector insertion with and without ITR, and unclassified reads.

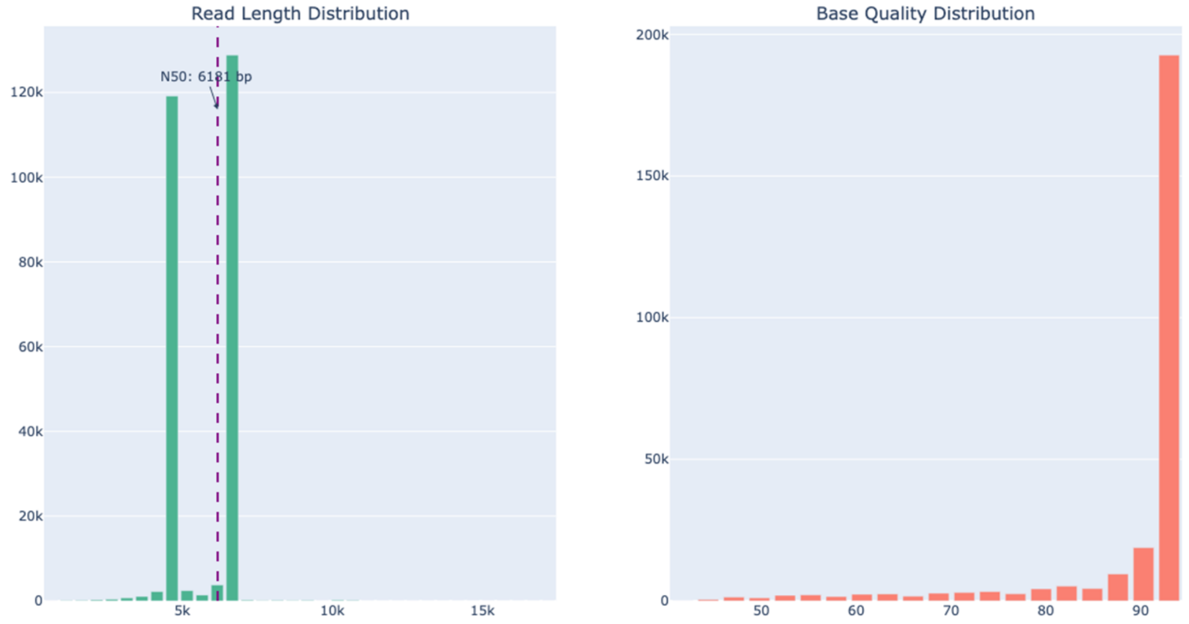


**Figure S2.** Representative read length and base quality distribution histograms from quality-control assessment.

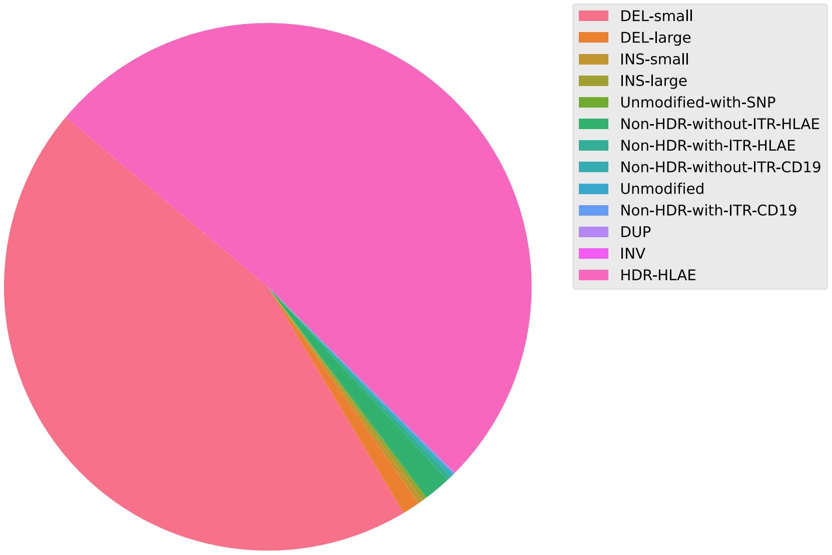


**Figure S3.** Representative pie chart showing the distribution of variant categories in a gene-edited sample.

| #Sample | Unmodified | DEL-small | INS-small | DEL-large | INS-large | .. |
|---|---|---|---|---|---|---|
| Sample A | 215 | 111705 | 748 | 2654 | 552 | 236 |
| Sample B | 133 | 99395 | 608 | 2630 | 351 | 230 |
| ... | 123 | 95051 | 620 | 2544 | 723 | 152 |

**Figure S4.** Example merged count table summarizing variant frequencies across multiple samples.

**Table S1. Accuracy of read classification in simulated datasets**

| Group | Simulation Category | Classification results | | | | | Accuracy(%) |
|---|---|---|---|---|---|---|---|
| | | INDEL Small* | SNP | HDR | Non-HDR w/ ITR | Unmodified | |
| 1 | HDR | | | 20000 | | | 100.0 |
| 2 | Full ITR forward | | | | 20000 | | 100.0 |
| 3 | Full ITR rev_com | | | | 20000 | | 100.0 |
| 4 | WT | 472 | 9 | | | 19519 | 97.60 |
| 5 | 1-side ITR left forward | | | | 20000 | | 100.0 |
| 6 | 1-side ITR left rev | | | | 20000 | | 100.0 |
| 7 | 1-side ITR right forward | | | | 20000 | | 100.0 |
| 8 | 1-side ITR right rev | | | 1 | 19999 | | 100.0 |
| 9 | 2-side ITR 50% del | | | | 20000 | | 100.0 |
| 10 | 2-side ITR 50% del rev | | | | 20000 | | 100.0 |
| 11 | 1-side ITR 50% del left | | | | 20000 | | 100.0 |
| 12 | 1-side ITR 50% del right | | | | 20000 | | 100.0 |
| 13 | 2-side ITR SNP | | | | 20000 | | 100.0 |
| 14 | 1-side ITR SNP right | | | | 20000 | | 100.0 |
| 15 | 1-side ITR SNP left | | | | 20000 | | 100.0 |

* INDEL Small includes reads classified as DEL-small and INS-small. SNP, HDR, Non-HDR w/ ITR, and Unmodified correspond to the original ALPINE categories. Blank cells indicate zero reads. Other ALPINE categories (e.g., DEL-large and INS-large) are not shown because they were zero across all simulation groups. Each simulation group contains 20,000 reads.

**Table S2. Read classification of two-target edits – Transgene A**

| Sample | Unmod /SNP | Indel <50bp | DEL >50bp | INS >50bp | HDR - A | Non-HDR | | | | INV | DUP | Total Reads |
|---|---|---|---|---|---|---|---|---|---|---|---|---|
| | | | | | | w/o ITR-A | w/ ITR-A | w/o ITR-B | w/ ITR-B | | | |
| Sample_1 | 798 | 31,744 | 4,882 | 1,504 | 115,539 | 7,303 | 2,122 | 947 | 851 | 48 | 1 | 165,739 |
| Sample_2 | 905 | 39,643 | 5,561 | 2,223 | 96,453 | 7,948 | 2,470 | 866 | 1,656 | 130 | 2 | 157,857 |
| Sample_3 | 541 | 15,186 | 1,701 | 524 | 118,259 | 3,293 | 755 | 294 | 219 | 38 | 4 | 140,814 |
| Sample_4 | 524 | 19,231 | 2,504 | 727 | 114,047 | 3,931 | 905 | 305 | 272 | 36 | 3 | 142,485 |
| Sample_5 | 1,210 | 40,213 | 4,612 | 2,564 | 251,893 | 4,859 | 2,843 | 587 | 825 | 216 | 3 | 309,825 |

**Table S3. Read classification of two-target edits – Transgene B**

| Sample | Unmod /SNP | Indel <50bp | DEL >50bp | INS >50bp | HDR - B | Non-HDR | | | | INV | DUP | Total Reads |
|---|---|---|---|---|---|---|---|---|---|---|---|---|
| | | | | | | w/o ITR-A | w/ ITR-A | w/o ITR-B | w/ ITR-B | | | |
| Sample_1 | 312 | 86,098 | 3,693 | 929 | 118,721 | 4,639 | 1,546 | 374 | 238 | 12 | 38 | 216,600 |
| Sample_2 | 499 | 79,632 | 3,069 | 674 | 128,449 | 3,407 | 1,368 | 218 | 457 | 7 | 25 | 217,805 |
| Sample_3 | 363 | 100,003 | 2,630 | 351 | 118,834 | 3,840 | 594 | 333 | 185 | 7 | 45 | 227,185 |
| Sample_4 | 451 | 112,453 | 2,654 | 552 | 128,817 | 3,549 | 652 | 385 | 209 | 11 | 28 | 249,761 |
| Sample_5 | 275 | 95,671 | 2,544 | 723 | 94,644 | 2,482 | 954 | 252 | 355 | 16 | 66 | 197,982 |

**Table S4. Summary of gene edits**

| | TransGene A | | | | TransGene B | | | |
|---|---|---|---|---|---|---|---|---|
| **Sample** | **HDR KI** | **Indels** | **Non-HDR** | **Other** | **HDR KI** | **Indels** | **Non-HDR** | **Other** |
| **Sample_1** | 69.711% | 23.006% | 6.771% | 0.511% | 54.811% | 41.884% | 3.138% | 0.167% |
| **Sample_2** | 61.102% | 30.044% | 8.197% | 0.657% | 58.974% | 38.280% | 2.502% | 0.244% |
| **Sample_3** | 83.982% | 12.365% | 3.239% | 0.414% | 52.307% | 45.330% | 2.180% | 0.183% |
| **Sample_4** | 80.041% | 15.764% | 3.799% | 0.395% | 51.576% | 46.308% | 1.920% | 0.196% |
| **Sample_5** | 81.302% | 15.295% | 2.942% | 0.461% | 47.804% | 49.973% | 2.042% | 0.180% |

HDR KI includes HDR category. Indels includes DEL-small, DEL-large, INS-small, and INS-large categories. Non-HDR includes Non-HDR-w/o-ITR-A, Non-HDR-w/o-ITR-B, Non-HDR-w/-ITR-A, and Non-HDR-w/-ITR-B categories. Other includes INV, DUP, SNP, and Unmodified categories.

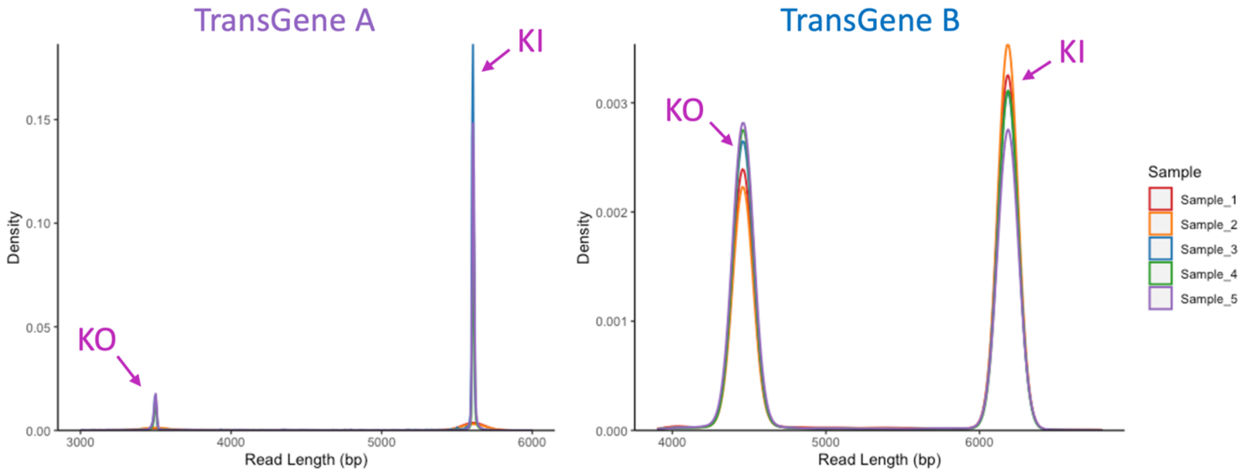


**Figure S5. Read length distributions of five samples.** Peaks for knock-out (KO) and knock-in (KI) are marked with arrowheads in the histogram. KO peak includes reads from Indels and Other categories. KI peak includes reads from HDR KI and Non-HDR categories.
